# Development of a BM7G((TKO/hCD46/hCD55/hTHBD/hEPCR) Donor Pig with Endogenous Promoter-Driven Transgenes for Xenotransplantation

**DOI:** 10.64898/2026.03.10.710833

**Authors:** Cong Xia, Meng Lian, Bingxiu Ma, Hongliang Yu, Renquan Zhang, Lijia Wen, Xueliang Wang, Yu Zhao, Zhen Ouyang, Yinghua Ye, Xiner Feng, Han Wu, Liangxue Lai

## Abstract

Xenotransplantation offers a potential solution to the organ shortage crisis. Multi-gene modification of pigs, such as knockout of three carbohydrate antigen-related genes and expression of immunoprotective proteins, can significantly improve xenograft survival. However, existing strategies face challenges: transposon-based transgenesis may lead to unstable expression, while exogenous promoters used in site-specific integration are susceptible to epigenetic silencing, hindering long-term stable expression. Therefore, developing a donor pig model capable of sustained multi-gene expression is critical. To address this, CRISPR-Cas9 was used to knockout three major glycan antigen genes to eliminate hyperacute rejection. Subsequently, four human protective genes were site-specifically integrated into the porcine *Rosa26* safe-harbor locus, with expression driven by the endogenous *Rosa26* promoter and the *THBD* core promoter for long-term stable and tissue-specific expression, and the selection marker was removed using Cre/loxP. Results showed complete absence of the three glycan antigens in BM7G pigs, while the four protective proteins were stably expressed in vascular endothelial cells and major organs. Among them, hCD55 and hCD46 were widely expressed, while hTHBD and hEPCR showed vascular-specific expression. *In-vitro* assays confirmed that BM7G porcine endothelial cells significantly reduced human antibody binding, effectively inhibited complement-dependent cytotoxicity, and decreased thrombin-antithrombin complex formation. In conclusion, by combining xenoantigen knockout with endogenous promoter-driven expression of multiple human protective genes, a seven-gene modified pig model with low immunogenicity and synergistic protective function was successfully constructed, providing an important donor resource for xenotransplantation preclinical research.

## 1 Introduction

Genetically modified pigs represent the most promising source of organs for xenotransplantation to address the global shortage of donor organs^[1]^. Key strategies include genetic modifications to mitigate xenogeneic immune rejection^[2]^, such as the elimination of three major carbohydrate xenoantigens (αGal, Neu5Gc, and Sda) via simultaneous inactivation of *GGTA1*, *CMAH*, and *β4GalNT2* genes, which significantly reduces pre-existing antibodies mediated rejection^[3]^. What’s more, expression of human complement regulatory proteins (e.g., hCD46, hCD55) and coagulation regulatory proteins (e.g., thrombomodulin, endothelial protein C receptor) in pigs effectively mitigates xenogeneic immune rejection and prolongs the survival time of grafts after transplantation^[4]^. To date, various genetically modified pigs have been developed^[4–7]^, achieving significant progress in pig-to-non-human primates heart, kidney and islet transplantation^[8–10]^, pig-to-brain-dead human, kidney^[11]^, liver^[12]^, and lung transplantation^[13]^, and even clinical studies^[14, 15]^.

Currently, multi-gene modifications in pigs primarily rely on two strategies: transgenic approaches and site-specific knock-in, each presenting distinct challenges. Although the PiggyBac transposon-based transgenic technology enables efficient integration, random insertion may lead to unstable expression and copy number variations in offspring^[16]^, affecting the reliable inheritance of phenotypes. Site-specific knock-in strategies based on homology-directed repair (HDR)^[17]^ or recombinase-mediated cassette exchange (RMCE)^[18, 19]^ allow precise integration of the target gene into predetermined genomic loci (e.g., safe harbor sites^[20]^), which significantly improves the genetic stability of the transgene. However, even with targeted integration, the exogenous promoters used may still be susceptible to host epigenetic silencing mechanisms^[21, 22]^, leading to gradual attenuation of transgene expression and making it difficult to maintain long-term, stable expression levels *in-vivo*^[23]^.

Using endogenous promoters to drive transgenes is an effective strategy for achieving long-term expression^[24]^. According to Li et al., the endogenous promoter of the porcine *Rosa26* (*pRosa26*) safe harbor locus exhibits high expression activity in various organs (e.g., liver, kidney, heart)^[25]^. Similarly, the porcine thrombomodulin (*pTHBD*) promoter efficiently drives the expression of *pTHBD* in porcine endothelial cell lines, making it suitable for driving human THBD expression, with stable inheritance to offspring^[26]^. In this study, we report the generation of a seven-gene-modified Bama miniature pig model. Three glycoantigen-related genes were knocked out, while four human protective genes were inserted into the *pRosa26* locus. Specifically, the complement regulators hCD55 and hCD46 are expressed under the control of the endogenous *pRosa26* promoter, whereas the coagulation regulators hTHBD and hEPCR are regulated by the *pTHBD* core promoter. This strategy effectively avoids the common issue of epigenetic silencing associated with exogenous promoters by utilizing endogenous promoters, thereby providing a reliable solution for generating multi-genetically modified pig models with stable and long-term transgene expression.

## 2 Methods

### 2.1 Pig welfare

Animal studies were conducted with the approval of the Institutional Animal Care and Use Committee (IACUC) of GIBH. All animals were managed under standard husbandry practices, and daily health monitoring to ensure their well-being. Following the collection of blood samples, ear tissue, and other specimens, gene-edited Bama minipigs were humanely euthanized via intravenous injection of an overdose of pentobarbital, ensuring rapid and stress-free death. Organs and tissues, such as kidneys and aortas, were subsequently harvested. Upon completion of all experiments, animal carcasses were incinerated at a licensed facility in compliance with biosafety and environmental protection regulations.

### 2.2 Generation of guide RNA plasmid for triple knockout

The sequences of the target genes *GGTA1*, *β4GALNT2*, and *CMAH* were download from the NCBI database. Guide RNAs (gRNAs) were designed using the online software (https://zlab.bio/guide-design-resources). gRNA-specific primers were synthesized by IGEbio Genomics Co., Ltd. And the sequences are show as Table.S1 Subsequently, the three gRNAs were subcloned into lentiCRISPR v2 (Addgene, cat. #52961) plasmid.

### 2.3 Assembly of *Rosa26* HDR constructs

For Rosa26 site-specific insertion, a homologous recombination repair (HDR) template vector was constructed. A 1.2 kb left homology arm and a 1 kb right homology arm were amplified from genomic DNA isolated from Bama miniature pigs. The human coding DNA sequences (CDSs) of human CD55 (hCD55), human CD46 (hCD46), human thrombomodulin (hTHBD), and human endothelial protein C receptor (hEPCR) were synthesized by IGEbio Genomics Co., Ltd. Two polycistronic transcription units were arranged in opposite orientations: one contains a viral splice acceptor (SA), a promoterless hCD55 CDS, and a hCD46 CDS, which are linked by a viral 2A peptide sequence; the other, in the inverted orientation, comprises an hTHBD CDS and hEPCR CDS, driven by the *pTHBD* core promoter (amplified from Bama miniature pig genomic DNA).

### 2.4 Transfection and selection of positive cell colonies

PFFs were cultured and electroporated using the Neon system (1350 V, 30 ms, 1 pulse) with plasmid. After recovery, cells were plated at low density and selected with puromycin (1000 ng/mL) for 14 days. Cell colonies were expanded and genotyped by PCR and sequencing using primers flanking the target loci (*GGTA1*, *β4GALNT2*, *CMAH*) and the *Rosa26* homologous arms.

### 2.5 Production of clone piglets

Gene-edited cells were cloned into pigs by SCNT, and cloning was performed as previously described^[27–29]^. Porcine oocytes were enucleated, injected with donor cells, and activated by electrofusion. Reconstructed embryos were cultured briefly and then surgically transferred into synchronized surrogate sows. Cloned piglets were ultimately delivered via cesarean section. The cloned piglets were delivered by cesarean section in DPF facility

### 2.6 Isolation of porcine peripheral blood mononuclear cells (PBMCs)

Fresh anticoagulated whole blood was collected and mixed with an equal volume of PBS. An equal volume of Ficoll-Paque (GE Healthcare, Cat. #17-1440-03) was added into a centrifuge tube, and the diluted blood was slowly layered onto the surface of the separation medium while maintaining a clear interface. After centrifugation at 400×g for 30 minutes, the solution separated into distinct layers. The PBMC fraction layer was carefully aspirated with a pipette and transferred to a new centrifuge tube. The harvested cells were washed with PBS, subjected to erythrocyte lysis, and then counted.

### 2.7 Flow cytometric phenotyping of PBMCs

PBMCs from wild-type (WT) and BM7G pigs were isolated using Ficoll-Paque Plus (GE Healthcare, Cat. #17-1440-03) according to the manufacturer’s protocol. Each sample (100,000 cells per sample) stained with primary and secondary antibodies. aGal was stained by FITC conjugated Isolectin B4 (IB4, SIGMA, Cat. #L2895, 1:100 dilution). β4GalNT2 phenotype was carried out using Fluorescein Dolichos Biflorus Agglutinin (DBA, Cat. Vector Laboratories, FL-1031, 1:50 dilution). The expression level of Neu5Gc was assessed by chicken anti-Neu5Gc antibody (BioLegend, Cat. #146901, 1:300 dilution), the secondary antibody was Alexa Fluor568 goat anti-chicken (Abcam, Cat. #ab96947, 1:1000 dilution), and the chicken IgY Isotype was used as negative control (BioLegend, Cat. #402101). Samples were washed and analyzed on a flow cytometer (CytoFLEX, Beckman). The data were collected and analyzed using FlowJo software (Flowjo_v10.8).

### 2.8 Antibody binding assay

Human sera were obtained from de-identified remnant clinical laboratory samples with approval. PBMCs from WT, TKO, and BM7G pigs were isolated using Ficoll-Paque Plus (GE Healthcare, Cat. #17-1440-03) according to the manufacturer’s protocol. Each sample (100,000 cells per sample) were incubated with inactivated 40% human serum for 30 min at 4°C, then washed three times with PBS and blocked with 5% BSA. Subsequently, the cells were stained with goat anti-human IgG Alexa Fluor 488 (Invitrogen, Cat. #A11013; 1:200 dilution) and goat anti-human IgM Alexa Fluor 647 (Invitrogen, Cat. #A21249; 1:200 dilution) for 30 min at room temperature. After three additional washes with PBS, flow cytometric analysis was performed using a CytoFLEX flow cytometer, and data were analyzed with FlowJo software (Flowjo_v10.8).

### 2.9 Detection of transgenic protein expression levels in tissues by western blot

Heart, liver, and kidney tissues from BM7G and TKO pigs were homogenized, and proteins were extracted. Protein concentrations were determined by BCA assay. Western blot was performed using standard electrophoresis, transfer, blocking, and ECL detection procedures. The expression of hCD55, hCD46, hTHBD, and hEPCR was detected using the following primary antibodies: Anti-CD55 Antibody (Bioster, Cat. #BM5281, 1:1000 dilution), Anti-CD46 antibody (Abcam, Cat. #AB108307, 1:1000 dilution), Anti-Thrombomodulin antibody (Abcam, Cat. #AB6980, 1:1000 dilution), and Anti-EPCR/CD201 antibody (Abcam, Cat. #AB236517, 1:1000 dilution). The corresponding secondary antibodies used were Goat Anti-Mouse IgG H&L (HRP) (Abcam, Cat. #ab6789, 1:5000 dilution) and Goat Anti-Rabbit IgG H&L (HRP) (Abcam, Cat. #ab6789, 1:5000 dilution).

### 2.10 Detection of surface antigens and protective proteins on porcine cells by IHC

Tissue sections were deparaffinized, underwent antigen retrieval, and were blocked with 3% BSA. After overnight primary antibody incubation at 4°C and PBS washes, sections were incubated with fluorescent secondary antibody for 1h in the dark, followed by DAPI nuclear staining. Images were captured using fluorescence microscopy. Following final washes, the sections were observed and images were captured using a fluorescence microscope. The primary antibodies used were as describe above, same as used in Western Blotting. The corresponding secondary antibodies used were Goat Anti-Mouse IgG H&L (Alexa Fluor® 488) (Abcam, Cat. #ab150113, 1:1000 dilution) and Goat Anti-Rabbit IgG H&L (Alexa Fluor® 594) (Abcam, Cat. #ab150080, 1:1000 dilution).

### 2.11 Isolation of porcine vascular endothelial cells (PECs)

Following euthanasia of pigs, the aortas were aseptically isolated and transferred into 150-mm sterile culture dishes. The specimens were rinsed repeatedly with sterile PBS, and redundant connective and adipose tissues surrounding the vessels were completely removed. The aortas were gently everted to fully expose the endothelial surfaces, followed by digestion with collagenase type Ⅳ (Sigma-Aldrich, Cat. #G5138) at 38℃ for 2 h. After the digestion was terminated with stop buffer, porcine vascular endothelial cells (PECs) were gently scraped 8 times in a single direction using a sterile cell scraper. All washing suspensions were collected, and the cells were harvested by centrifugation and cryopreserved separately. The isolated cells were stained with PECAM-1 Alexa Fluor® 488-conjugated Antibody (R&D Systems, Cat. #FAB33871G-100ug) to determine their purity.

### 2.12 Complement-dependent cytotoxicity (CDC) assay

WT, TKO, and BM7G endothelial cells (100,000 cells per sample) were incubated with 40% human serum at room temperature for 30 min. After incubation, the cells were stained with propidium iodide (PI). After 3 minutes, the percentage of PI-positive cells was detected using a CytoFLEX flow cytometer.

### 2.13 TAT formation assay

WT, TKO, and BM7G endothelial cells were seeded in 24-well plates at a density of 300,000 cells per well. After 24 hours of culture, the cells were incubated with fresh human whole blood collected in anti-coagulant tubes. Following incubation at 37°C with gentle shaking for 1 hour, the plasma was separated. The concentration of the thrombin-antithrombin (TAT) complex in the plasma was measured using a Human Thrombin–Antithrombin Complex ELISA Kit (ab108907, Cat. # Abcam).

### 2.14 Statistics analysis

Statistical analysis was performed using GraphPad Prism 8 software. A two-tailed Student’s t-test or ANOVA was used for comparisons, and statistical significance was defined as a P value of < 0.05

## 3 Results

### 3.1 Production of BM7G donor pigs

The engineering workflow was presented in Fig. 1A. Firstly, Cas9 expression plasmid together with sgRNAs targeting the *GGTA1*, *CMAH*, and *β4GALNT2* genes was delivered into wild-type (WT) Bama PFF cells via electroporation (Table. S1). Triple-gene knockout (TKO) cell clones were obtained and used for somatic cell nuclear transfer (SCNT) and embryo transfer. A pregnant surrogate was sacrificed at day 33 after the embryo transfer. Twenty fetuses were collected to isolate PFF cells, genotyped results indicated 10 of these fetuses were biallelic TKO (numbers: 9#-14#, 16#, 17#, 18#, 20#) (Fig. 1B) (Table. S2). Then, a HDR knock-in vector targeting the *pRosa26* locus was constructed, which contained two expression cassettes to enable the expression of four immune-protective proteins (hCD55, hCD46, hTHBD, and hEPCR) (Fig. 1C). This HDR donor vector, together with a Cas9 and *Rosa26*-sgRNA plasmid, were electroporated into TKO PFF cells. After selection with puromycin, positive *pRosa26* knock-in cell clones were obtained, which were confirmed to be correctly targeted by 5′- and 3′-arm PCR analysis (Fig. S1).

**Fig. 1.**
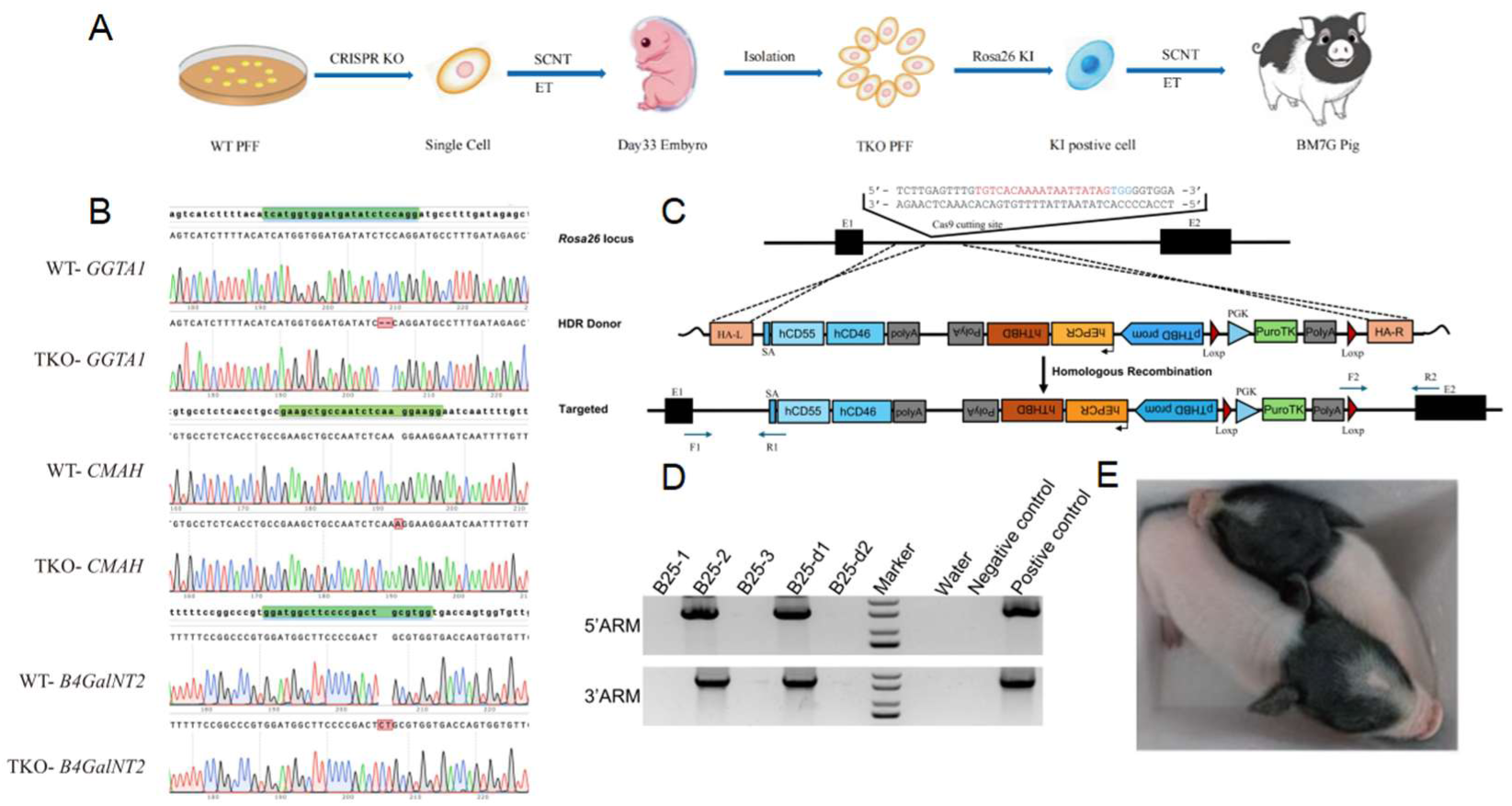
Generation of BM7G pigs for xenotransplantation. **A**. The workflow for generating BM7G pigs via continuous cloning and multiple rounds of gene editing. **B**. Sanger sequencing results of *GGTA1*, *CMAH*, and *β4GALNT2* in WT and Triple-KO positive fetuses. **C**. Strategy for targeted knock-in of a dual transgene expression cassette into the porcine *Rosa26* locus. **D**. Genotyping of piglets by PCR analysis. Piglets B25-2 and B25-d2 tested positive for both the 5’ (1382 bp) and 3’ (1614 bp) homology arms of the targeted *Rosa26* locus. **E**. BM7G-positive cloned piglets.

Subsequently, SCNT and embryo transfer were performed using these positive cells as nuclear donor. The 2 surrogate sows gave birth to 5 cloned piglets. Genotyping analysis confirmed that 2 of these piglets were successful knock-in at the *Rosa26* locus (Fig. 1D), with the genotype TKO/hCD55/hCD46/hTHBD/hEPCR, these piglets were termed as BM7G (Fig. 1E).

### 3.2 Cre/loxP-mediated excision of the selection marker

To obtain genetically modified cells without a selection marker, ear tissue was collected from BM7G donor pigs to isolate primary ear fibroblasts. Next, transient transfection of a Cre-expressing plasmid was performed to mediate the deletion of the selection marker between the two loxP sites (Fig. 2A). A total of 20 cell clones were picked and subjected to PCR genotyping. Gel electrophoresis results showed that, with the exception of clone 10#, the PGK-Puro selection cassette was successfully excised in the remaining 19 cell clones (Fig. 2B), representing an excision with high efficiency. Sanger sequencing further confirmed the precise removal of the selection marker, leaving only a single loxP (Fig. 2C). Then, the positive cells were used as nuclear donors for SCNT. A total of 676 embryos were transferred into 4 surrogates. Ultrasound examination one month later revealed that 3 of the sows were pregnant. The pregnant sows were maintained under specific pathogen-free conditions and finally delivered by cesarean section, yielding 12 piglets, 7 of which survived. PCR analysis of genomic DNA from the surviving piglets confirmed that all were derived from the above SCNT donor cells (Fig. 2D).

**Fig. 2.**
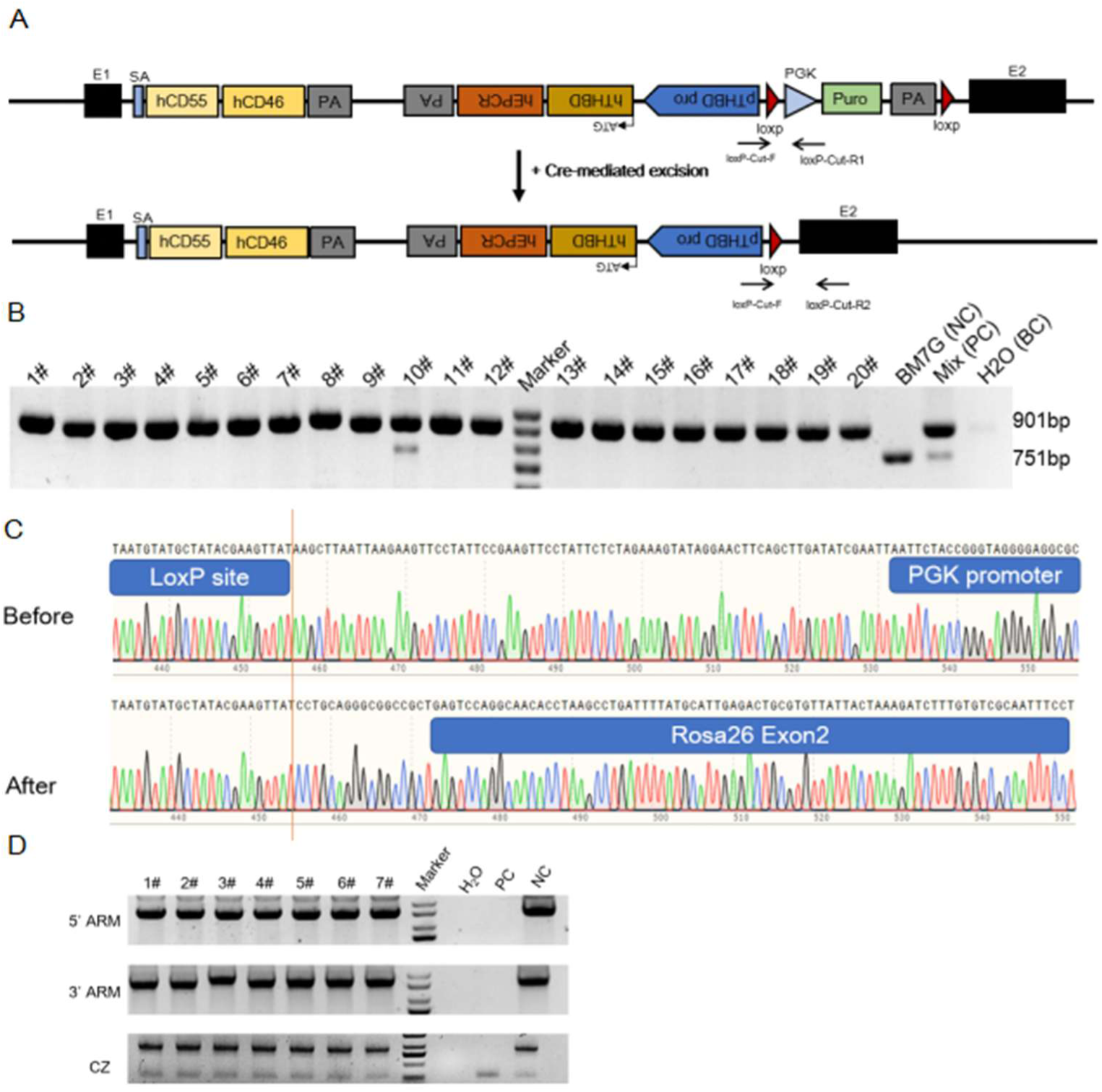
Cre-mediated deletion of selection marker. **A.** Strategy for the deletion of the Selection Marker. **B**. Genotyping of cell clones by PCR analysis. Untransfected cells (BM7G (NC)) served as the negative control, and a mix of cells collected during the experiment was used as the positive control. The specific bands at 901 bp and 751 bp confirm the precise deletion event. **C**. Comparison of Sanger sequencing chromatograms for the recombination locus. The lower panel (After) clearly shows the sequence junction after Cre recombinase treatment. **D**. Genotyping of Cloned Pig Offspring by PCR. Lanes 1-7: Seven different cloned pig samples; H₂O: no-template control (water); PC: Positive control (genomic DNA from BM7G pig); NC: Negative control (genomic DNA from wild-type pig).

### 3.3 Detection of protein expression at the cellular level

To examine whether three xeno-antigens (α-Gal, Sda, and Neu5Gc) were completely eliminated, peripheral blood mononuclear cells (PBMCs) were isolated from the WT and BM7G pigs. Subsequently, the PBMCs were stained with fluorescently labeled antibody, respectively. Flow cytometry analysis of PBMCs revealed complete absence of α-Gal, Sda, and Neu5Gc antigens in BM7G pigs (Fig. 3A), consistent with functional knockout of *GGTA1*, *CMAH*, and *β4galNT2* (Fig. 1F).

**Fig. 3.**
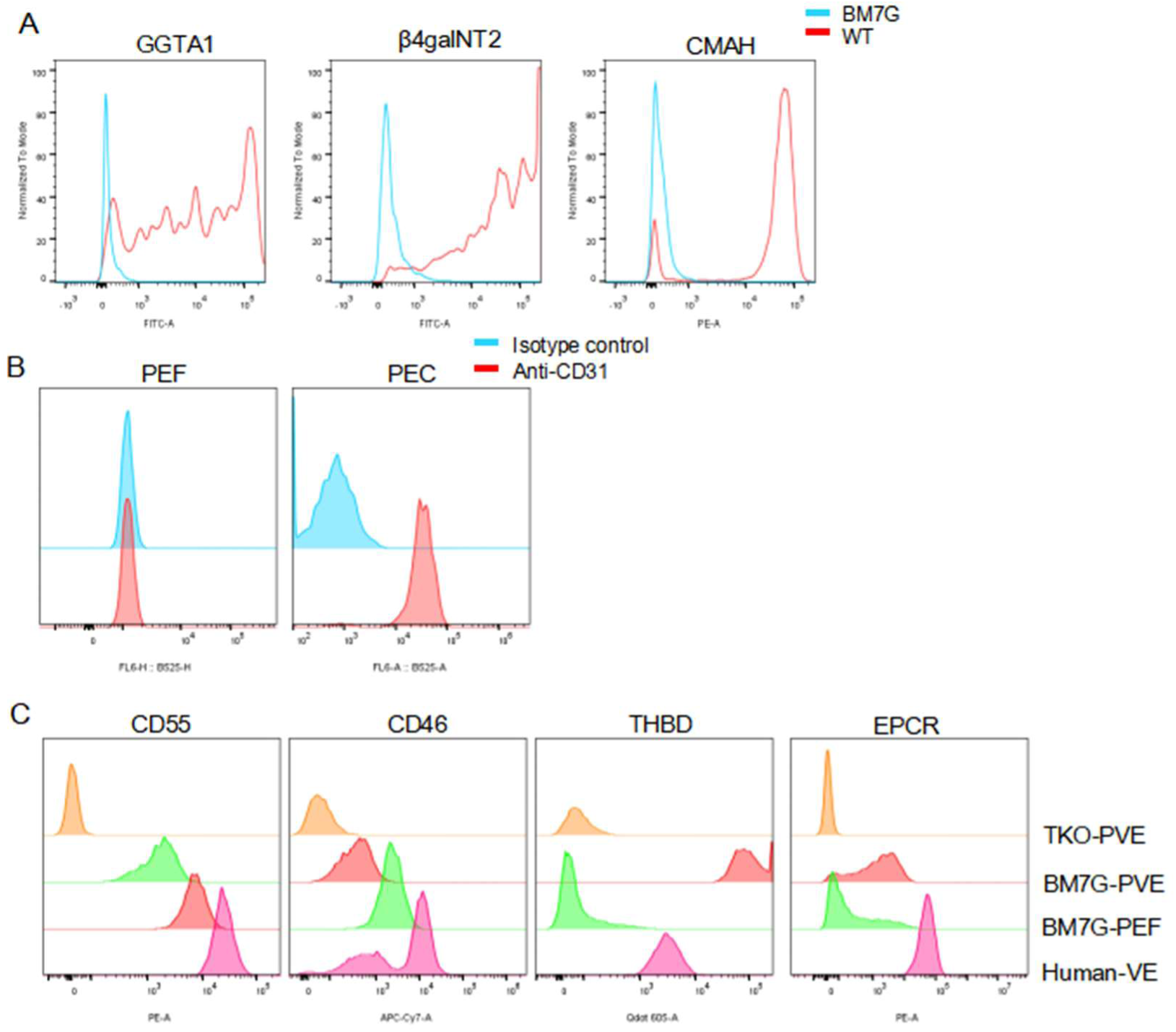
Detection of protein expression at the cellular level. **A.** Comparison of α-Gal (*GGTA1*), SDa (*β4GalNT2*), and Neu5Gc (*CMAH*) expression on PBMCs from BM7G (blue) and TKO (red) pigs. **B**. Detection of the endothelial marker CD31 on PEFs and PECs. **C**. Flow cytometric analysis of four immune-protective proteins in TKO-PEC, BM7G-PEC, BM7G-PEF, hVE.

While knocking out the three genes responsible for synthesizing these glycan epitopes, four immune-protective protein genes were integrated into the *Rosa26* locus. We assessed the expression patterns of these transgenes across distinct cell types of BM-7G pigs. TKO vascular endothelial cells (TKO-PEC), BM7G vascular endothelial cells (BM7G-PEC), and BM-7G ear fibroblasts (BM7G-PEF) were isolated and cultured (Fig. 3B). Then the transgene expression across different cell lines were analyzed by flow cytometry, with human vascular endothelial (VE) cells used as a control. Compared to TKO-PEC, BM7G-PEC exhibited significantly elevated expression of hCD55, hCD46, hTHBD, and hEPCR (Fig. 3C). In contrast, BM7G-PEF expressed only hCD55 and hCD46, consistent with the ubiquitous activity of the *Rosa26* promoter. This expression pattern further confirms the endothelium-specific activity of the *pTHBD* promoter. Notably, hCD55 and hCD46 expression levels in BM7G endothelial cells were comparable to those observed in human VE cells, whereas hTHBD expression was markedly higher.

### 3.4 Detection of protein expression at the organ level

To further verify the expression of these four proteins at the organ level, heart, liver, and kidney tissues were collected from the BM7G pig and performed RT-qPCR and Western blot analysis. At both the mRNA and protein levels, expression of the four immune-protective proteins was detected in all three tissues (Fig.4A, B). We also characterized the genetic modifications in heart, liver, and kidney tissues via immunofluorescence staining for the three glycan epitopes and four immune-protective proteins. Compared to the organ tissues from WT pig, α-Gal, Sda, and Neu5Gc antigens were completely negative in the heart, liver, and kidney tissues of BM-7G pigs (Fig. 4C, S2). All four transgenic proteins were successfully detected in the heart, liver, and kidney tissues of BM-7G pigs. The complement regulatory proteins, hCD55 and hCD46, were prominently expressed and widely distributed, while the coagulation regulatory proteins hTHBD and hEPCR were specifically localized to the vascular regions of these tissues (Fig. 4C, S2). This specific expression pattern indicates that the endogenous *pRosa26* promoter drives stable and ubiquitous expression of complement regulate proteins, whereas the *pTHBD* promoter exhibits strong endothelial specificity. Furthermore, immunohistochemical staining was performed to verify the expression of hCD55, hCD46, hTHBD, and hEPCR in the three organs of BM-7G pigs (Fig. 4D, S3). The results further confirmed stable expression of these four protective proteins at the organ level.

**Fig. 4.**
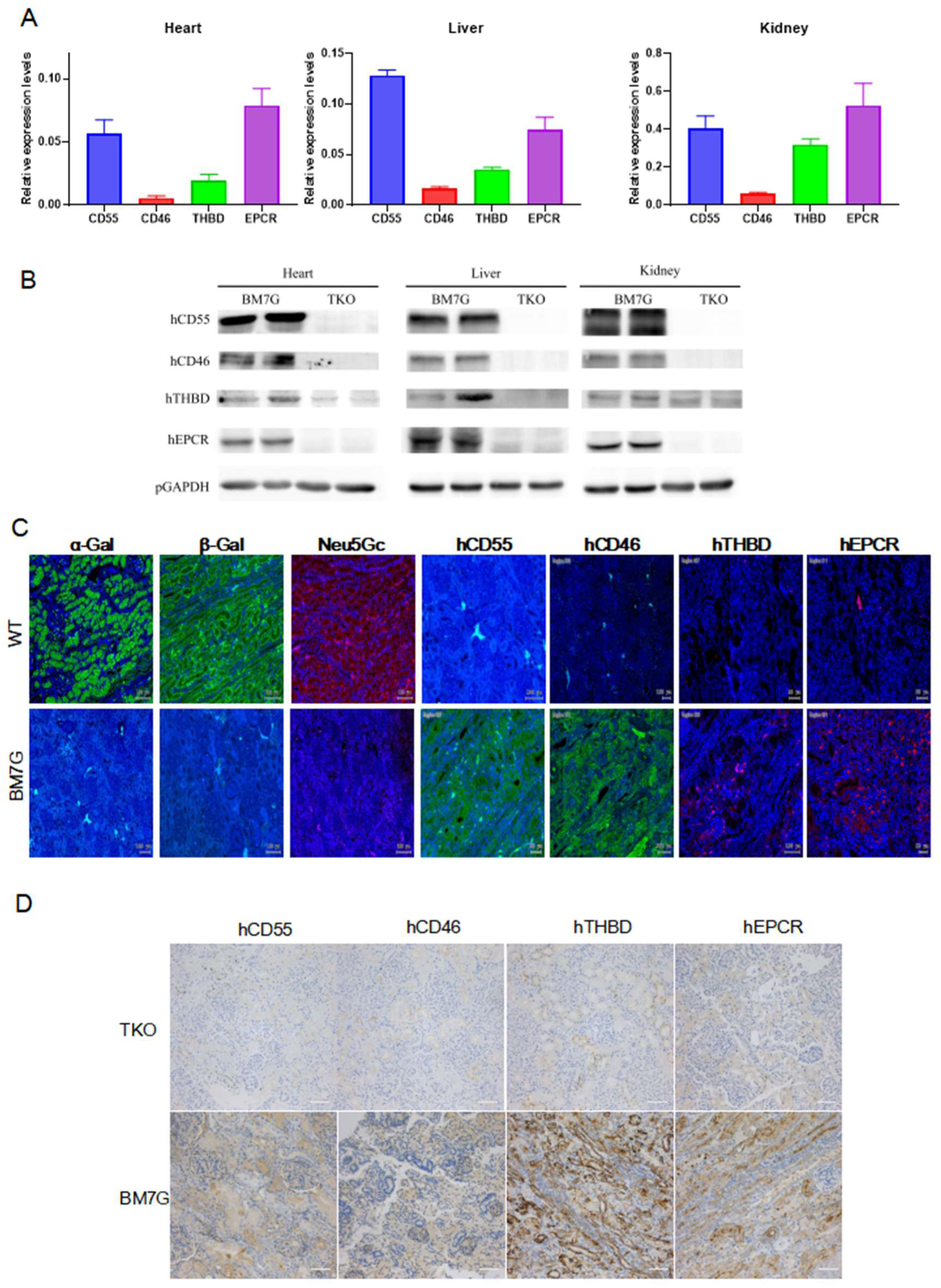
Detection of protein expression at the organ level. **A**. RT-qPCR analysis of immune-protective gene expression in heart, liver, and kidney. **B**. Western Blot analysis of immune-protective gene expression in heart, liver, and kidney. **C**. Immunofluorescence staining of kidney sections comparing WT and BM7G pigs. **D**. Immunohistochemical staining of human protective proteins in kidney sections comparing TKO and BM7G pigs.

### 3.5 Immunogenicity detection of BM7G

After confirming the negative status of the three glycan epitopes, the immunogenicity of BM-7G pig cells was evaluated by incubating PBMCs from WT, TKO, and BM-7G pigs with 40% heat-inactivated human serum. Significantly reduced binding of human IgM and IgG antibodies was observed in both BM-7G and TKO PBMCs compared to WT controls, indicating that knockout of the *GGTA1*, *β4GALNT2*, and *CMAH* genes greatly diminished xenograft immunogenicity. To further assess the function of the expressed complement regulatory proteins, CDC assays were performed using endothelial cells isolated from the pigs. The results showed that the TKO group significantly reduced complement-mediated cytotoxicity compared to the WT group. The positive rate of PI of the BM7G group were further reduced to nearly zero after background subtraction, demonstrating the functional activity of the expressed hCD55 and hCD46 (Fig.5C). Finally, the anti-coagulant functions of hTHBD and hEPCR were verified via a thrombin-antithrombin complex (TAT) assay, in which PEC from BM7G co-cultured with fresh human whole blood significantly reduced TAT formation compared to WT and TKO cells, indicating effective regulation of coagulation and potential alleviation of coagulation disorders in xenotransplantation (Fig.5D).

**Fig. 5.**
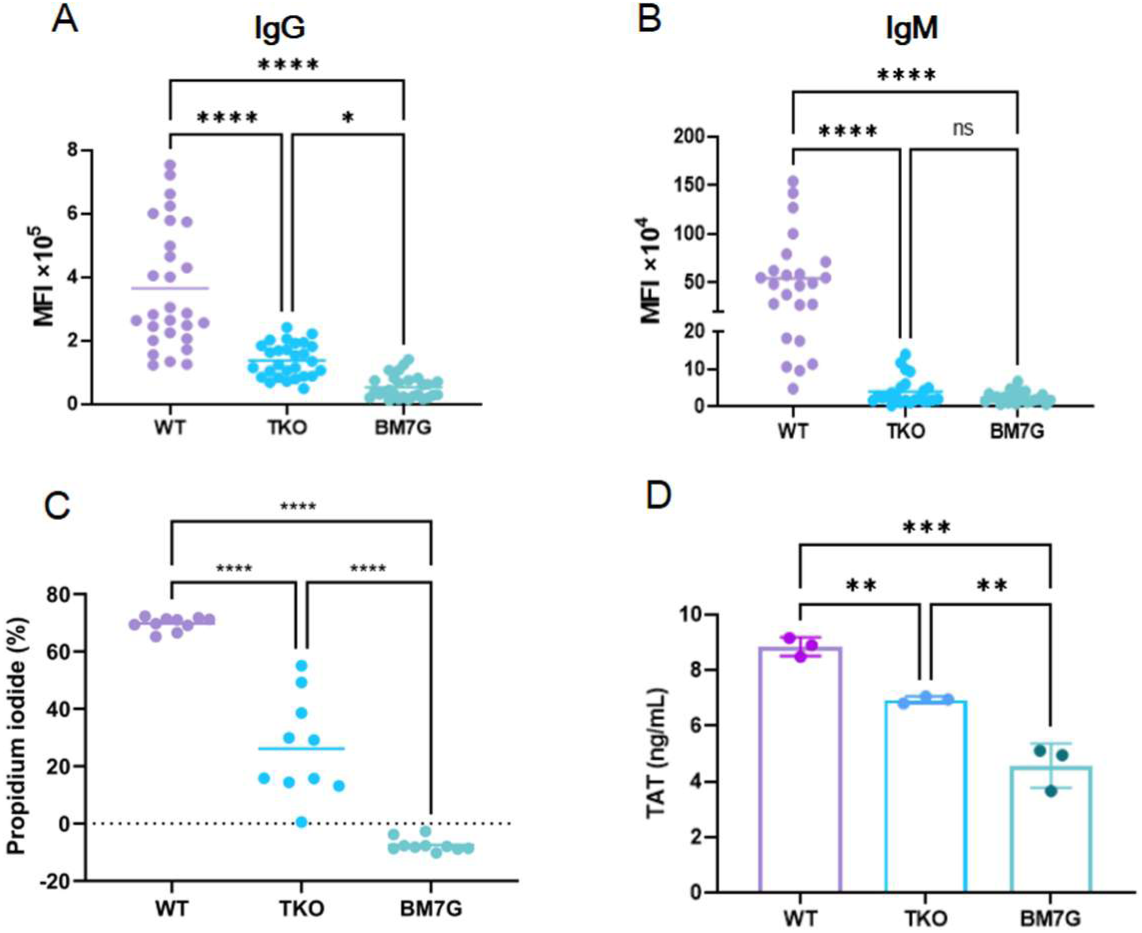
Immunogenicity detection of BM7G. Antibody binding assay: Flow cytometric analysis of human IgG (**A**) and lgM (**B**) binding to PBMCs isolated from WT, TKO, and BM7G pigs. **C**. Complement-dependent cytotoxicity (CDC) assay on porcine vascular endothelial cells. **D**. Thrombin-antithrombin (TAT) complex generation assay using porcine vascular endothelial cells. Statistical significance was determined by one-way ANOVA (****, p < 0.0001, ***, p < 0.001, **, p < 0.01). Data are shown as mean ± SD.

### 3.6 BM7G off-target assessment

To test whether off-targeting occurred in BM7G pig, potential off-target site of *GGTA1*, *CMAH*, *β4GalNT2* and *pRosa26* sgRNA were screened using Cas-OFFinder (http://www.rgenome.net/cas-offinder/)^[30]^. We amplified these off-target sites from BM7G pig genome by PCR and performed Sanger sequence. These results indicated that genome editing occurred in only the targeted region, and no off-target mutations were introduced at other untargeted sites (Table.1, Fig. S4).

**Table. 1.** BM7G off-target assessment.

| Gene | *GGTA1* | *β4GalNT2* | *CMAH* | *Rosa26* |
| --- | --- | --- | --- | --- |
| Number of predicted sites | 9 | 8 | 9 | 10 |
| Number of off-target sites | 0 | 0 | 0 | 0 |

## 4 Discussion

In this study, we successfully developed a novel seven-gene-modified xenotransplantation donor pig model. Its construction was based on two core genetic engineering strategies: first, the three major xenoantigen genes (*GGTA1*, *β4GalNT2*, and *CMAH*) were completely knocked out to eliminate hyperacute rejection; second, four human protective genes were site-specifically integrated into the *pRosa26* safe harbor locus. By utilizing endogenous promoters, these genes were stably expressed in various organs such as the heart, liver, and kidneys, driving the expression of human complement regulatory proteins (hCD55 and hCD46) and thrombomodulins (hTHBD and hEPCR) to simultaneously suppress complement activation and coagulation dysregulation^[31, 32]^. To evaluate their function, systematic *in-vitro* validation was performed. The results showed that PEC derived from BM7G pigs significantly alleviated pre-existing antibody-mediated immune rejection and effectively inhibited coagulation pathway activation, demonstrating that the multiple modifications exert synergistic protective effects at the cellular level.

Rosa26 is widely expressed in both embryonic and adult tissues^[33]^. Targeting genes to the Rosa26 locus represents an ideal method for creating transgenic animals with sustained and high-level transgene expression^[34, 35]^. Li et al^[25]^ successfully generated *pRosa26*-targeted pigs and observed widespread reporter gene expression at both fetal and adult stages, thereby validating the general applicability of this locus in pig models. However, in the field of xenotransplantation, when Fang et al^[36]^ integrated expression cassettes containing hCD55 and hCD47 genes driven by an IHK chimeric promoter into the *pRosa26* locus, an low level expression of these two proteins was detected at the erythrocyte or platelet level. This was likely due to epigenetically mediated transcriptional silencing or differences between human and porcine transcriptional mechanisms^[37, 38]^. Subsequent experiments in PK-15 cell lines further demonstrated that using the porcine endogenous *CD47* promoter could improve the expression levels of CD55 and CD47, suggesting that endogenous promoters play a crucial role in driving stable expression of exogenous immune-protective genes. Therefore, in this study, all human protective genes were site-specifically integrated into the *Rosa26* locus and driven by porcine endogenous promoters (*pRosa26* or *pTHBD* promoter) rather than traditional exogenous viral promoters. We confirmed that the *pRosa26* promoter enables broad transgene expression across multiple cell and organ types, while the *THBD* promoter exhibits a specific ability to drive expression in vascular endothelial cells.

The presence of a selection marker gene can limit subsequent genetic manipulations within the same individual, and the currently available types of selection markers are limited^[39]^. Therefore, its removal via the Cre/loxP system facilitates the future cultivation of donor pigs with more complex genetic modifications. At present, the elimination of selection markers has not received sufficient attention in the field of xenotransplantation. However, in the production of recombinant protein drugs, residual selection marker genes and their expression products may increase the immunogenicity risk. This approach not only eliminates associated safety and regulatory concerns but also lays the foundation for further model optimization to ultimately meet stringent clinical application requirements.

## 5 Conclusion

In a word, we successfully developed a BM7G pig model for xenotransplantation, utilizing endogenous promoters to drive the expression of human protective genes, which ensures more stable expression. The model demonstrated complete elimination of major xenoantigens and synergistic immune-protective functions in vitro. Further pig-to-non-human primate transplantation studies will be essential to evaluate the long-term safety, graft function, and immune rejuction of BM7G organs, thereby advancing their clinical translation.

## Author contributions

Cong Xia, Han Wu and Liangxue Lai conceived of the project. Cong Xia and Meng Lian designed the experiments. Cong Xia and Meng Lian completed the experimental design with help from Bingxiu Ma, Renquan Zhang, Xueliang Wang, Yu Zhao, and Zhen Ouyang. Cong Xia analyzed the data with help from Hongliang Yu, Bingxiu Ma, Xiner Feng and Lijia Wen. Cong Xia, Han Wu drafted the manuscript. Cong Xia, Han Wu and Liangxue Lai reviewed and edited the manuscript. Yinghua Ye was responsible for the procurement of experimental supplies. All authors read and approved the final manuscript.

## Compliance and ethics

The authors declare that they have no conflict of interest.

## Supplementary Figures

**Fig. S1.**
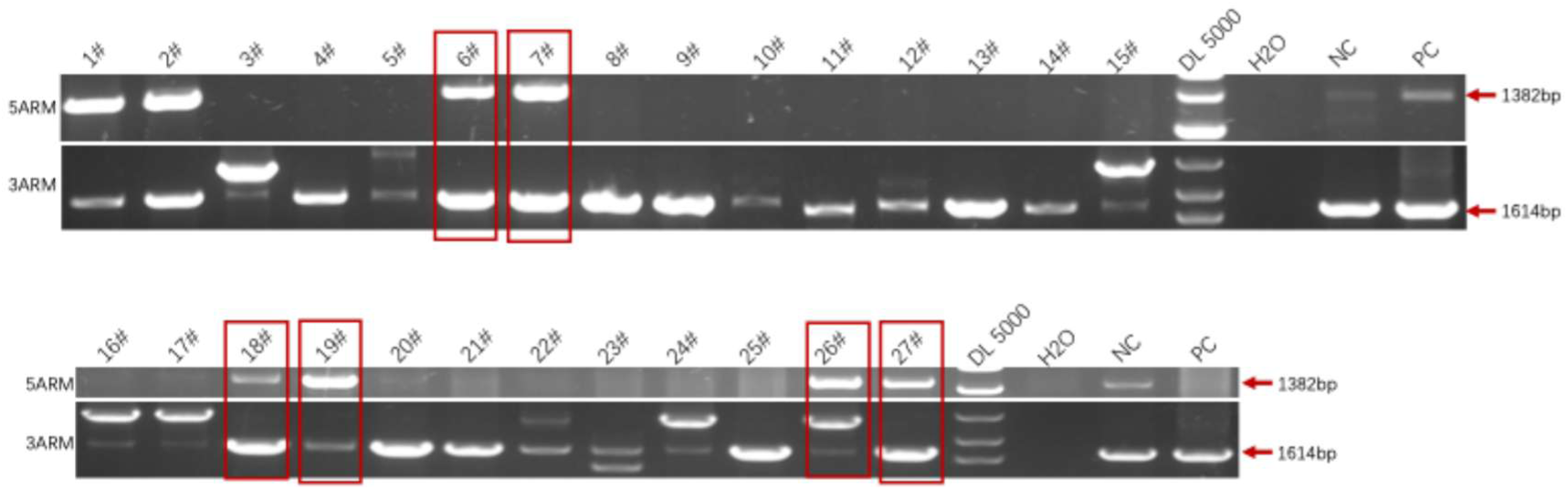
PCR results of the 5’ and 3’ ARM of the knock-in vector at the porcine *Rosa26* locus. The 5’ and 3’ arm bands are 1382 bp and 1614 bp, respectively. Cell clones 6#, 7#, 18#, 19#, 26#, and 27# are positive.

**Fig. S2.**
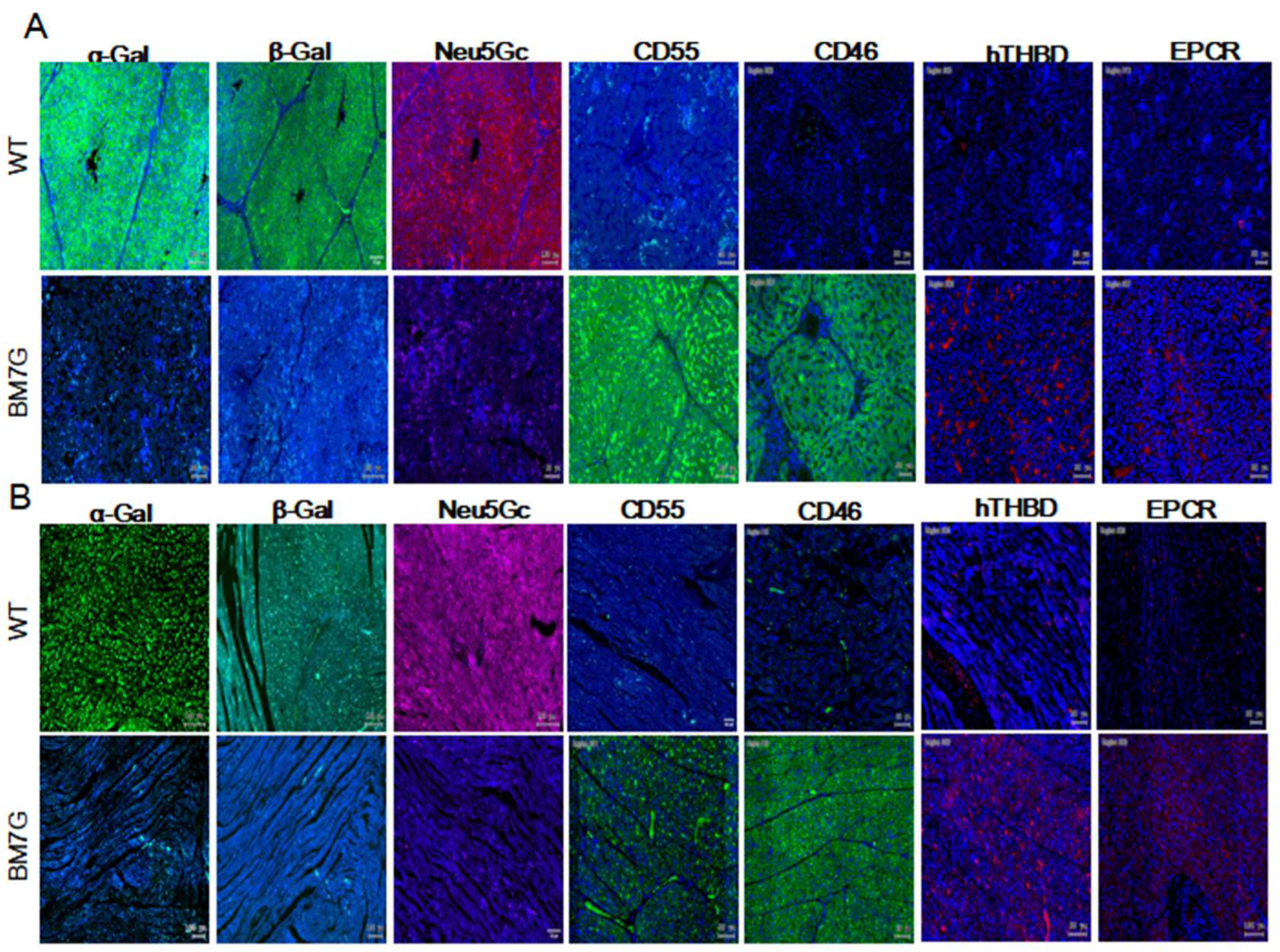
Immunofluorescence staining of heart (**A**) and liver (**B**) sections from wild-type (WT, upper panels) and BM7G (lower panels) pigs. The tissues were stained for the expression of three xeno-antigens (α-Gal, β-Gal, Neu5Gc) and four Immune compatibility proteins (hCD55, hCD46, hTHBD, hEPCR). The results demonstrate the absence of xeno-antigen expression and the successful detection of the human proteins in BM7G tissues. Experiments were independently repeated three times with consistent results. Scale bar = 100 μm.

**Fig. S3.**
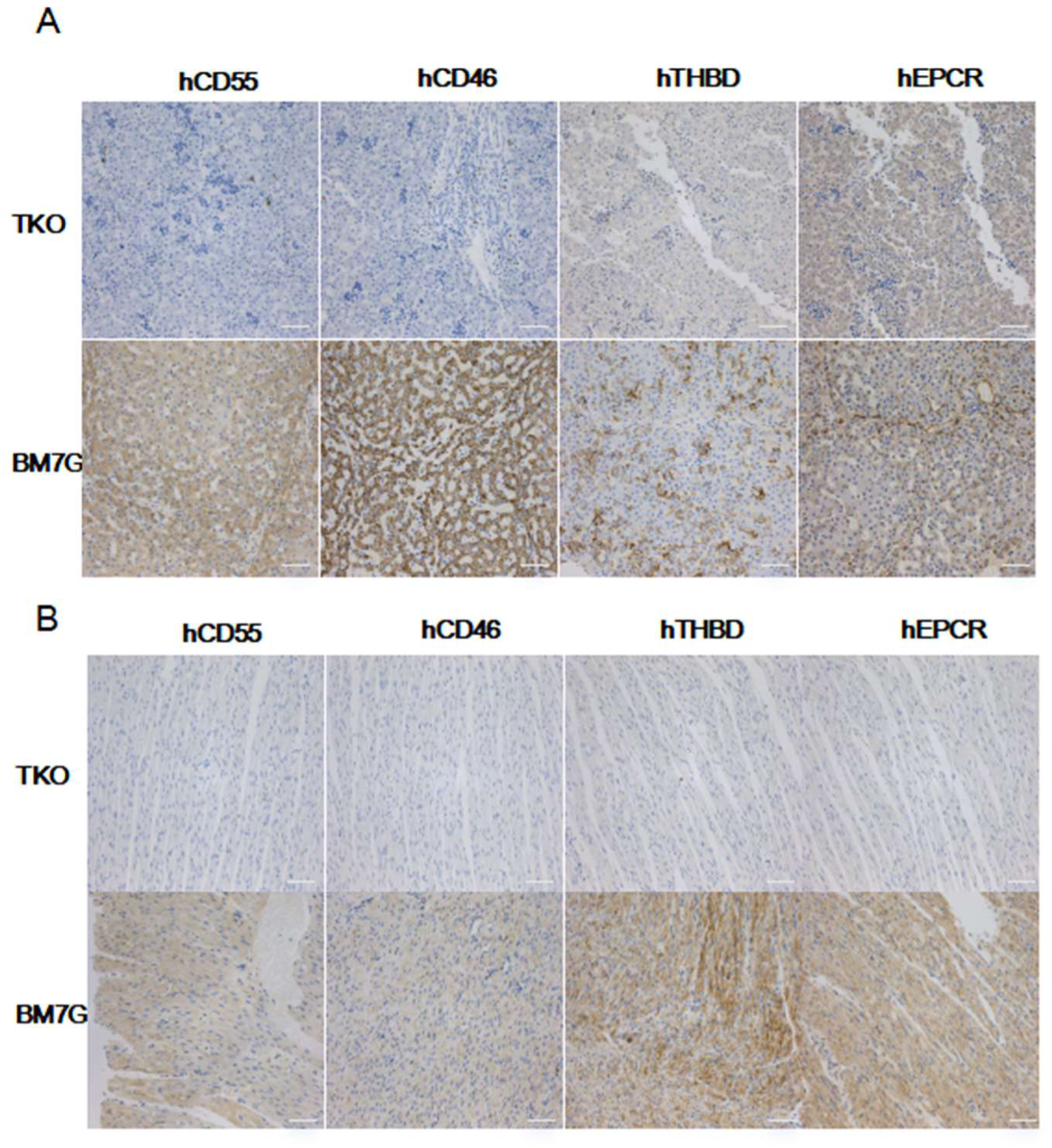
Immunohistochemical (IHC) analysis of heart (**A**) and liver (**B**) sections from wild-type (WT, upper panels) and BM7G (lower panels) pigs. The expression of four immune compatibility proteins (hCD55, hCD46, hTHBD, hEPCR) was detected. Scale bar = 100 μm.

**Fig. S4.**
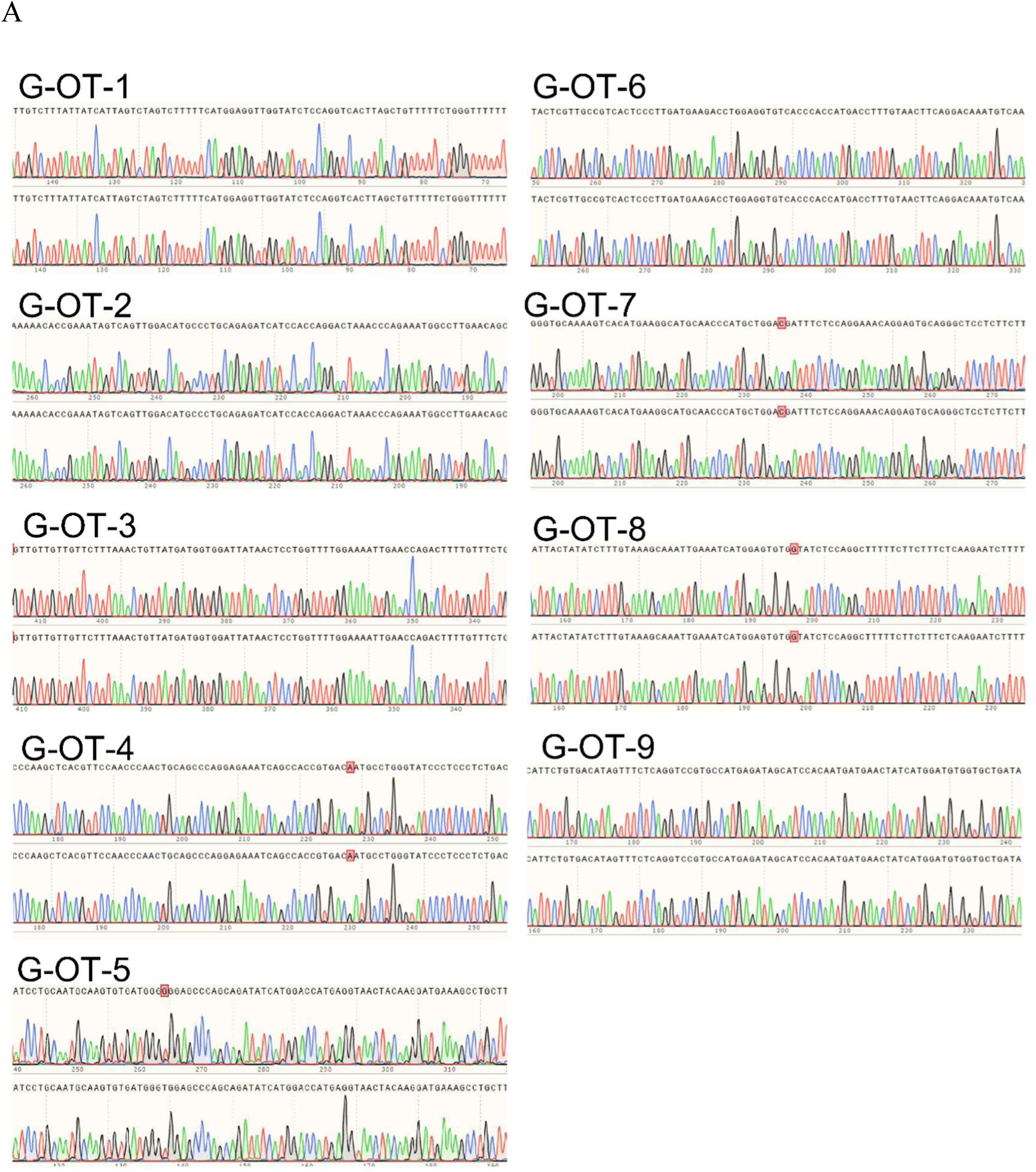
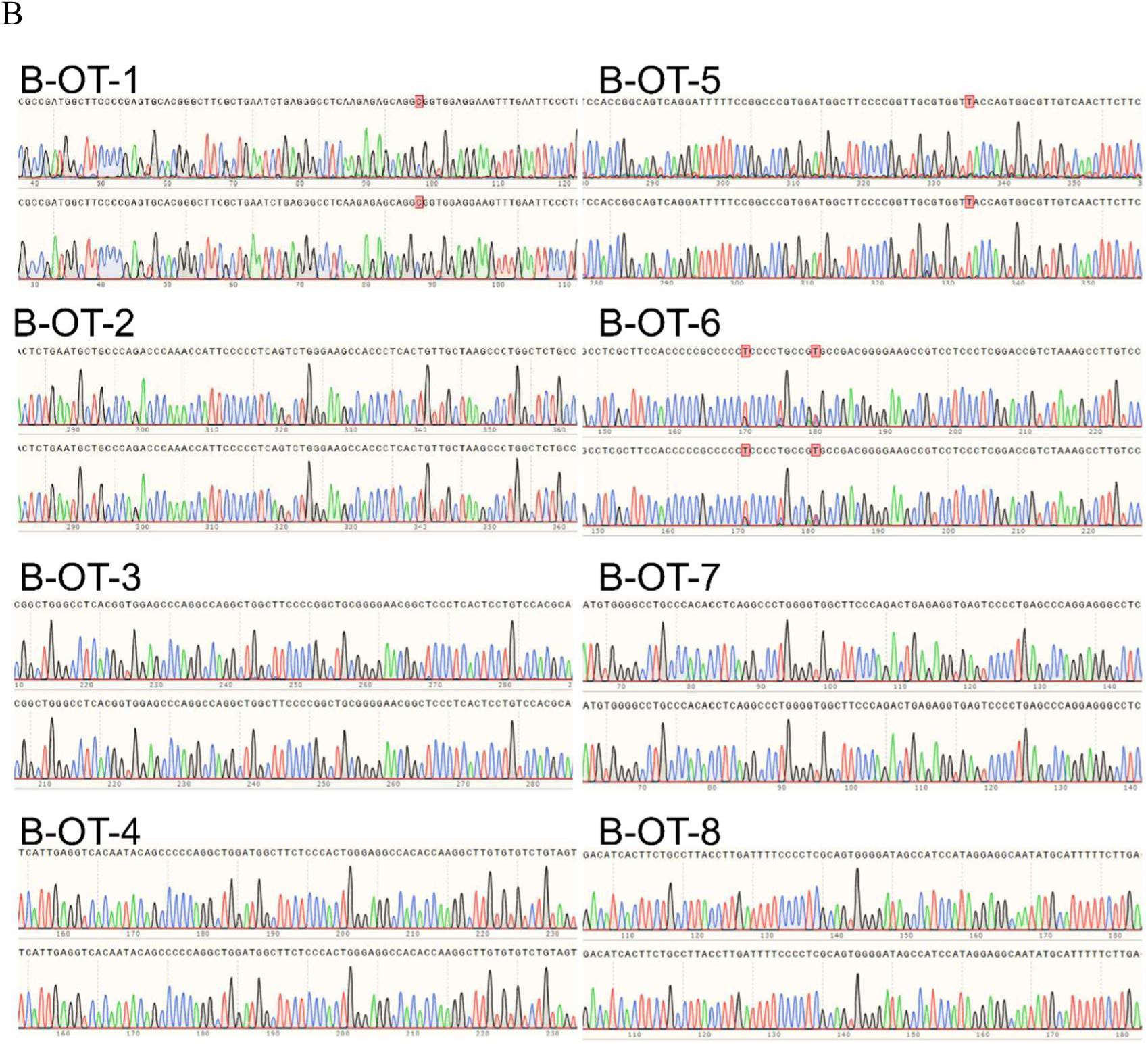
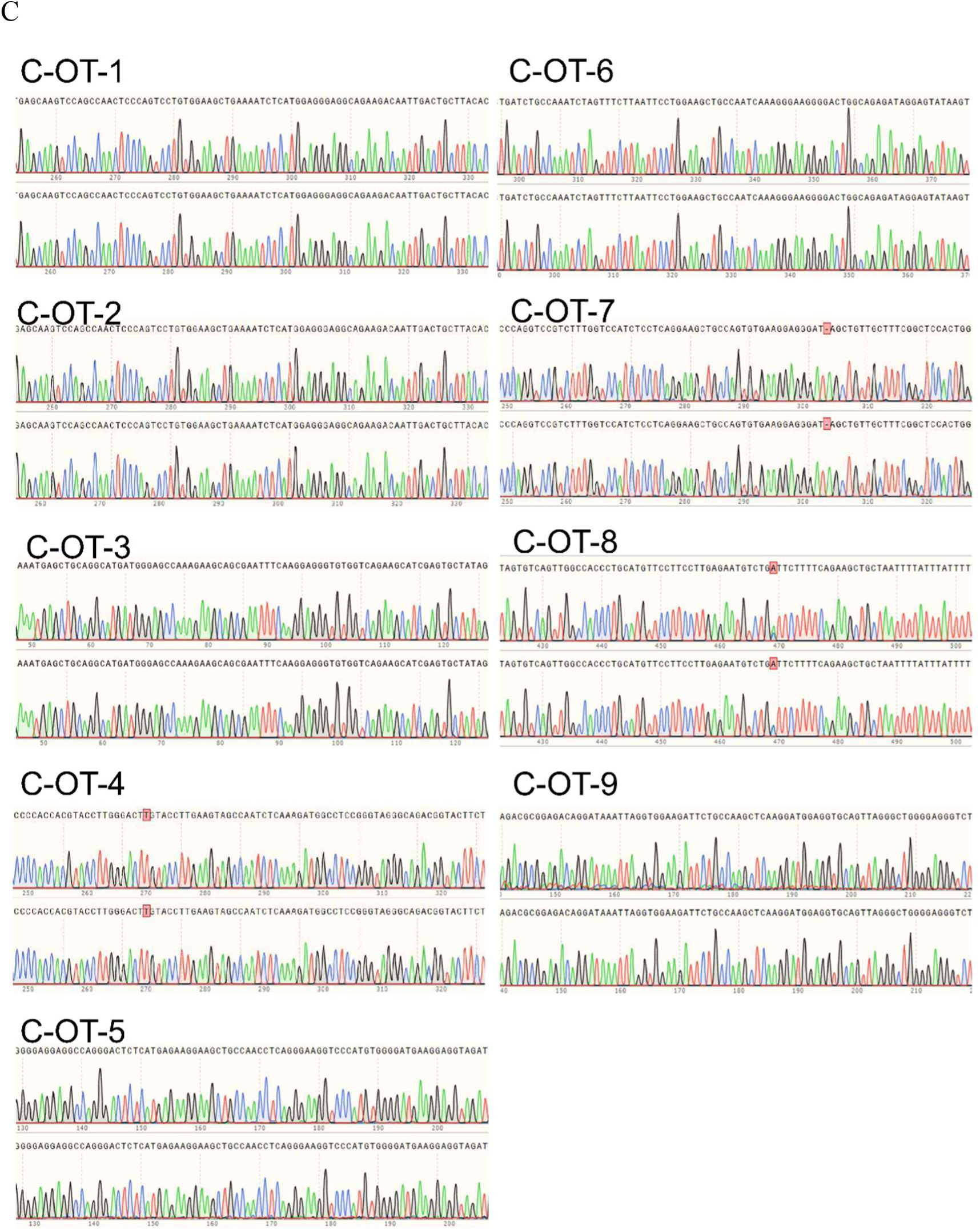
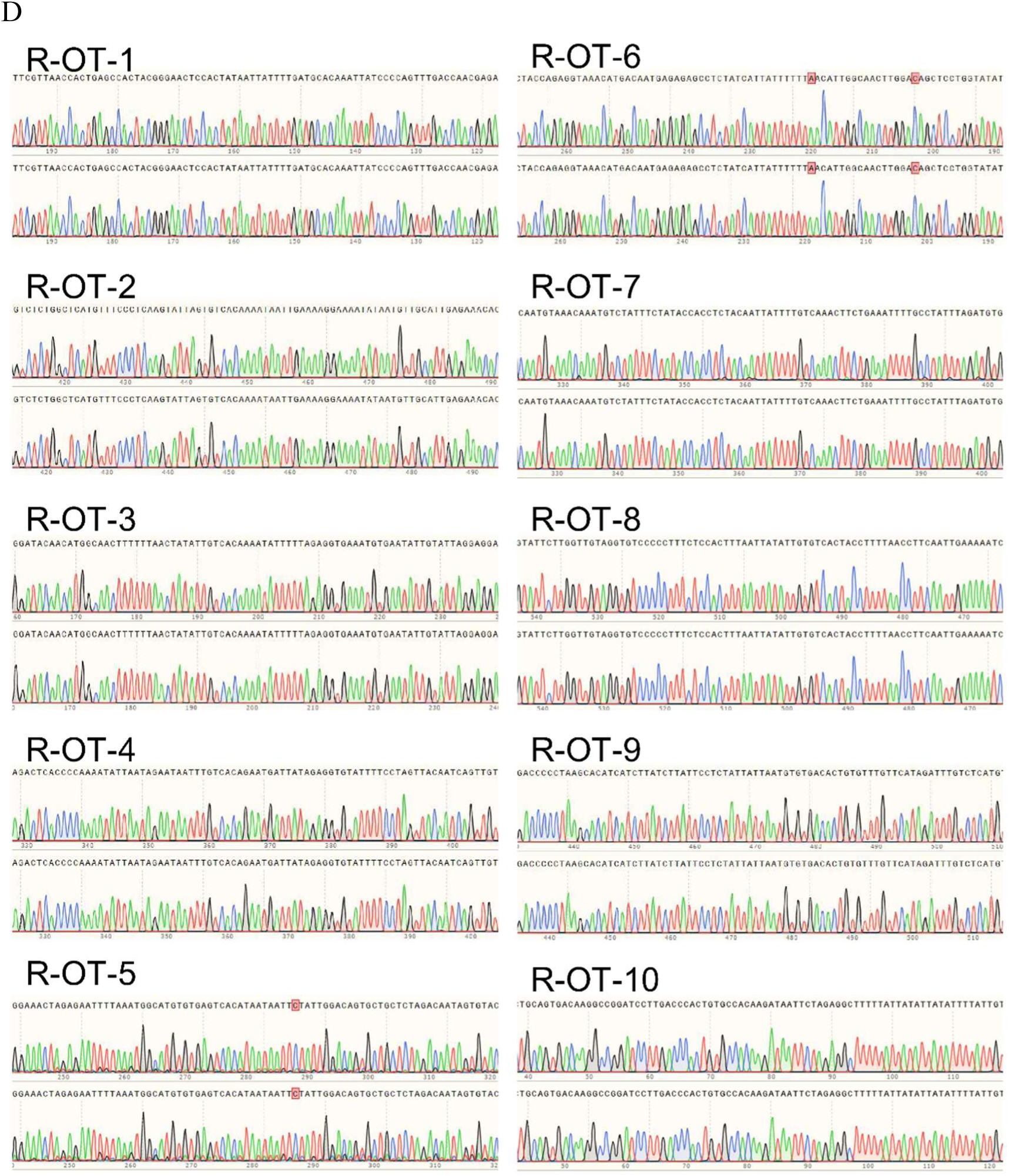
Off-Target Analysis for *GGTA1*, *CMAH*, and *B4GalNT2* at the *Rosa26* Locus: Sanger sequencing of WT (top) and TKO (bottom) cells shows identical sequences at predicted off-target sites for *GGTA1* **(A)**, *β4GalNT2* **(B)**, *CMAH* **(C)**, and *Rosa26* **(D)**, indicating no off-target editing.

